# Whole-brain drug distribution profiles of psychedelic drugs provide insights into rapid antidepressant action

**DOI:** 10.64898/2026.04.04.715307

**Authors:** Benjamin Hänisch, Tobias Kaufmann, Sofie L. Valk

## Abstract

Recent studies pioneered the use of classic hallucinogens as rapid-acting antidepressants (RAAD). To further understand the link between their neuromodulatory and antidepressant effects, we combine pharmacodynamic profiles of four classic hallucinogens and the anaesthetic Ketamine with receptor density distributions from both Positron Emission Tomography (PET) and layer-resoluted autoradiography studies to develop anatomical distribution profiles of drug action strengths giving a comparative measure how strong a drug would act in a region of interest. PET-based, we find high action strengths in association cortices for classic hallucinogens, which we contextualise anatomically using functional and cytoarchitectural measures. Autoradiography-based, we observe high action strengths in the supragranular layer and multimodal temporal areas. Finally, we show how Ketamine’s affinity to high-affinity subtypes of 5-HT2a and D2 receptors produce classic hallucinogen-like neuroanatomical trends. Through highlighting high RAAD action strengths in regions with emotion processing functionality, our results contribute to a mechanistic understanding of rapid antidepressant action.

## Introduction

Major depressive disorder is a common mental health condition that critically contributes to the global burden of disease [1]. Pharmacotherapy is an important pillar in its treatment, and conventional antidepressant drugs are empirically shown to be effective [2]. However, while their pharmacodynamic properties are known, their mechanisms of antidepressant action are still incompletely understood [3]. Conventional antidepressants generally require a daily dosage regimen to build a constant plasma level to be effective, and continual administration after clinical improvement is necessary to sustain symptom remission. Furthermore, they show a lag of weeks between treatment onset and antidepressant effect, while side effects can occur immediately after the first dosing. These factors negatively influence therapy adherence and complicate treatment [4]. In light of these issues, recent efforts have explored the antidepressant potential of psychedelic drugs as rapid-acting antidepressants (RAAD). RAAD differ from conventional antidepressants in three critical ways: they are not to be chronically administered, show antidepressant effects without a week-long time lag, and produce noticeable altered states of consciousness. Drug-induced psychedelic experiences may include alterations in visual and auditory sensory perception, hallucinations, derealisation and depersonalisation, of which the latter may be so intense that it can be experienced as complete ‘ego-dissolution’ [5]. There is ample evidence that experiencing these altered states of consciousness is instrumental in producing the antidepressant effect, as the quality of the psychedelic experience predicts long-term remission of depressive symptoms [6,7]. Furthermore, both the quality of the psychedelic experience as well as their antidepressant efficacy change in a dose-dependent manner [8,9]. Indeed, the nascent field of psychedelic-assisted psychotherapy aims to further exploit therapeutically beneficial elements of psychedelic experiences, which are hypothesised to reduce pathological avoidance and facilitate the processing of highly emotionally distressing events [10].

To improve upon the development and clinical use of RAAD, the neural basis of psychedelic experiences and their antidepressant effects need to be better understood. Although multiple explanatory hypotheses incorporating neuroanatomical, neuromodulatory and circuitory elements exist, they offer different and sometimes conflicting explanations for the drugs’ effects. For example, an early mechanistic account proposed a disruption of cortico-striato-thalamic loops that result in impaired thalamic gating [11], while a more recent theory with a strong focus on subcortical structures proposes that canonical cortical network architecture is attenuated through decreased cortico-claustro-cortical coupling [12]. Furthermore, while theories on the brain’s role as a computational inference machine agree in identifying prior beliefs as a promising target for psychedelic-induced alteration, they disagree on the exact nature of the perturbation necessary to cause psychedelic experiences. On the one hand, the prominent Relaxed Belief under Psychedelics (REBUS) model posits that psychedelics decrease the weighting of high-level beliefs and thereby indirectly increase the importance of sensory information in inference processes [13]. The theory similarly posits that inflexible high-level beliefs residing in the Default Mode Network (DMN) [14] contribute to depressive etiology. On the other hand, the Strong Prior model claims that psychedelics’ effects like hallucinations are best described by an increased weighting of high-level priors [15], without making any definitive anatomical claims.

Explanatory accounts for antidepressant and psychedelic action of RAAD are further complicated by the apparent difference in pharmacodynamic properties of classic hallucinogens and the dissociative anesthetic drug Ketamine, the only RAAD approved for clinical use by major regulatory bodies to date. Ketamine primarily acts as an NMDA receptor antagonist, while the antidepressant and psychedelic effects of classic hallucinogens are thought to be mediated through strong agonism at the metabotropic serotonin receptor 5-HT2a, which also takes center stage in the REBUS model [13]. Incidentally, there is evidence suggesting that Ketamine similarly acts at high-affinity subtypes of 5-HT2a and D2 receptors [16]. Additionally, since psychedelics also act strongly at other metabotropic receptors next to 5-HT2a, most prominently among them the 5-HT1a serotonin receptor, the exact pharmacodynamic basis of rapid antidepressant action remains unclear [17].

Summarised, a better understanding of the functional macro-anatomical and neuromodulatory targets of RAAD are needed to improve mechanistic explanations of rapid antidepressant action. Given the clear links between the brain’s functional organisation and the organisation of its neurotransmitter receptor and transporter landscape [18–20], this study looks to exploit neuroanatomical knowledge to further understand functional properties of rapid antidepressant action. To this end, we synthesise the well-established pharmacodynamic properties of classic hallucinogens and Ketamine [16,21] with neurotransmitter receptor density maps derived from whole-brain Positron Emission Tomography (PET) and post mortem autoradiography studies [19,22,23] to calculate anatomical drug distribution profiles across the whole cortex and select subcortical structures, with additional layer-resoluted distribution profiles in available cortical areas. As this approach relies on no specific hypothesis which ligand-receptor interaction mediates rapid antidepressant effects, it allows for the generation of unbiased anatomical patterns that can be interpreted and contextualised against the broader literature of functional anatomy [24–26].

## Results

In order to investigate neuroanatomical properties of rapid-acting antidepressants, we combined pharmacodynamic profiles, which can be found in **Table 1**, and the brain’s underlying neurochemical architecture, to calculate anatomical drug distribution profiles. Briefly summarised, the affinities were used to weight the influence of their respective receptors in summing the underlying receptor densities to a unit-less value per brain region that describes a pharmacodynamic-informed measure of importance, or strength, of that brain region in the drug’s action relative to all other brain regions (for a detailed description, see the **Methods** section).

**Table 1.** Ki values of RAAD substances given in nanoM, extracted from [16,21].

|  | **Psilocin** | **DMT** | **LSD** | **Mescaline** | **Ketamine** | **Ketamine**<br>**(high affinity)** |
| --- | --- | --- | --- | --- | --- | --- |
| **NMDA** | 100000 | 100000 | 100000 | 100000 | 500 | 500 |
| **D2** | 3700 | 3000 | 25 | 100000 | 100000 | 500 |
| **5-HT2a** | 49 | 237 | 4 | 6300 | 100000 | 15000 |
| **5-HT1a** | 123 | 75 | 3 | 4600 | 100000 | 100000 |
| **alpha1** | 6700 | 1300 | 670 | 100000 | 100000 | 100000 |
| **alpha2** | 2100 | 2100 | 12 | 1400 | 100000 | 100000 |
| **D1** | 100000 | 6000 | 310 | 100000 | 100000 | 100000 |

### PET decoding of classic hallucinogens

We first calculated drug distribution profiles of classic hallucinogens from parcellated PET-derived neurotransmitter density maps of the human cortex and projected them upon the cortical surface (**Fig. 1A**). Generally, drug action strengths of psychedelics were lower in the paracentral lobule and occipital cortices, and stronger in the temporal lobe, with especially high values in the inferior temporal gyrus, the temporal pole and the insular cortex. To further contextualize the drug distribution profiles with existing normative neuroanatomical measures, we quantified their distribution across functional networks derived from resting-state fMRI, and two different brain parcellations based on cytoarchitectural characteristics – first, a sensory-fugal gradient that spans different levels of cognitive processing complexity across four different cytoarchitectural classes [25], and second, laminar architecture-derived cytoarchitectural classes from the Structural Model of cortical hierarchies [24] (**Fig. 1B**). The decoding analysis showed similar trends in Mescaline, DMT and LSD. Here, functional network decoding revealed the lowest average drug action strength values in the somato-motor network and the highest in the limbic network [26]. In the cytoarchitectural classes of the sensory-fugal gradient, drug action strengths were lowest in idiotypic cortices and highest in paralimbic cortices, with uni- and heteromodal cortices showing similar middling drug action strengths. Laminar-derived cytoarchitectural decoding showed strongest drug action strengths in dysgranular cortices, and a steady decrease of drug action strengths with increasing levels of myelination across the eulaminate cortices. Psilocin showed highest drug action strengths in the DMN, heteromodal cortices and least myelinated Eulaminate cortices (Eu I).

**Figure 1.**
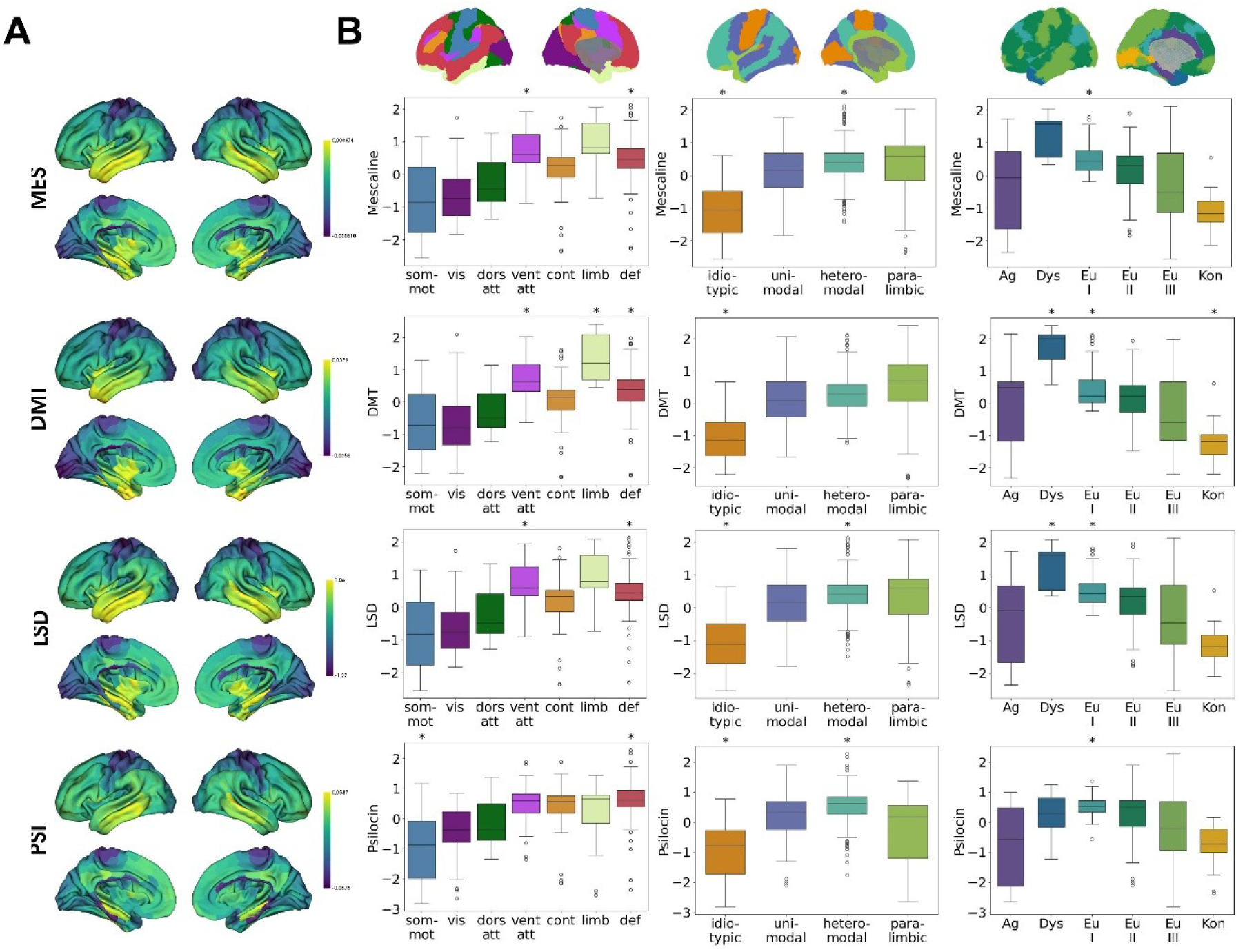
PET-derived cortical drug distribution profiles of classic hallucinogens. **A)** Cortical projections of drug distribution profiles of Mescaline (MES), LSD, DMT and Psilocin (PSI). **B)** Neuroanatomical decoding of classic hallucinogens’ drug distribution profiles, measured as the distribution of their values through three neuroanatomical modes. Projections of the respective modes are displayed on the cortical surface above the boxplots. *Left:* Distribution across seven functional networks (som-mot: Somato-motor; vis: visual; dors att: dorsal attention; vent att: ventral attention; cont: control; limb: limbic; def: default mode). *Middle:* Distribution across four cytoarchitectural classes. *Right:* Distribution across six layering-derived cortical types (Ag: Agranular; Dys: Dysgranular; Eu I: Eulaminate I; Eu II: Eulaminate II; Eu III: Eulaminate III; Kon: Koniocortex). Asterisks indicate significant alignment to the respective region as determined by spin permutation testing at significant levels of p < 0.05.

### Layer-wise decoding of select cortical areas

Our previous results revealed that classic hallucinogens show heterogeneous drug distribution profiles across the cortical surface. As a next step, we looked to also assess vertical cortical anatomical differentiations, to gain insights into how classic hallucinogens’ action might be distributed in the layered anatomy of the cortical sheet, since cortical layering is an important determinant of multiple histological features, including the brain’s neurotransmitter receptor architecture [27]. To this end, we calculated drug distribution profiles using autoradiographic measures of neurotransmitter receptor concentrations across a set of 44 cortical areas defined by features of cyto- or receptor architecture (**Table 2**, **Figure 2A**) which differentiate between infragranular, granular and supragranular cortical layers. Drug distribution profiles across cortical areas were generally similar between classic hallucinogens. Strikingly, drug action strengths were highest in the supragranular layer across cortical areas and across classic hallucinogens drugs (**Figure 2B**). Regarding the distribution across cortical areas and substances, drug action strengths generally peaked in the multimodal temporal areas 21 and 36 and were at their nadir in areas 4 and 6, as determined by z-scores.

**Figure 2.**
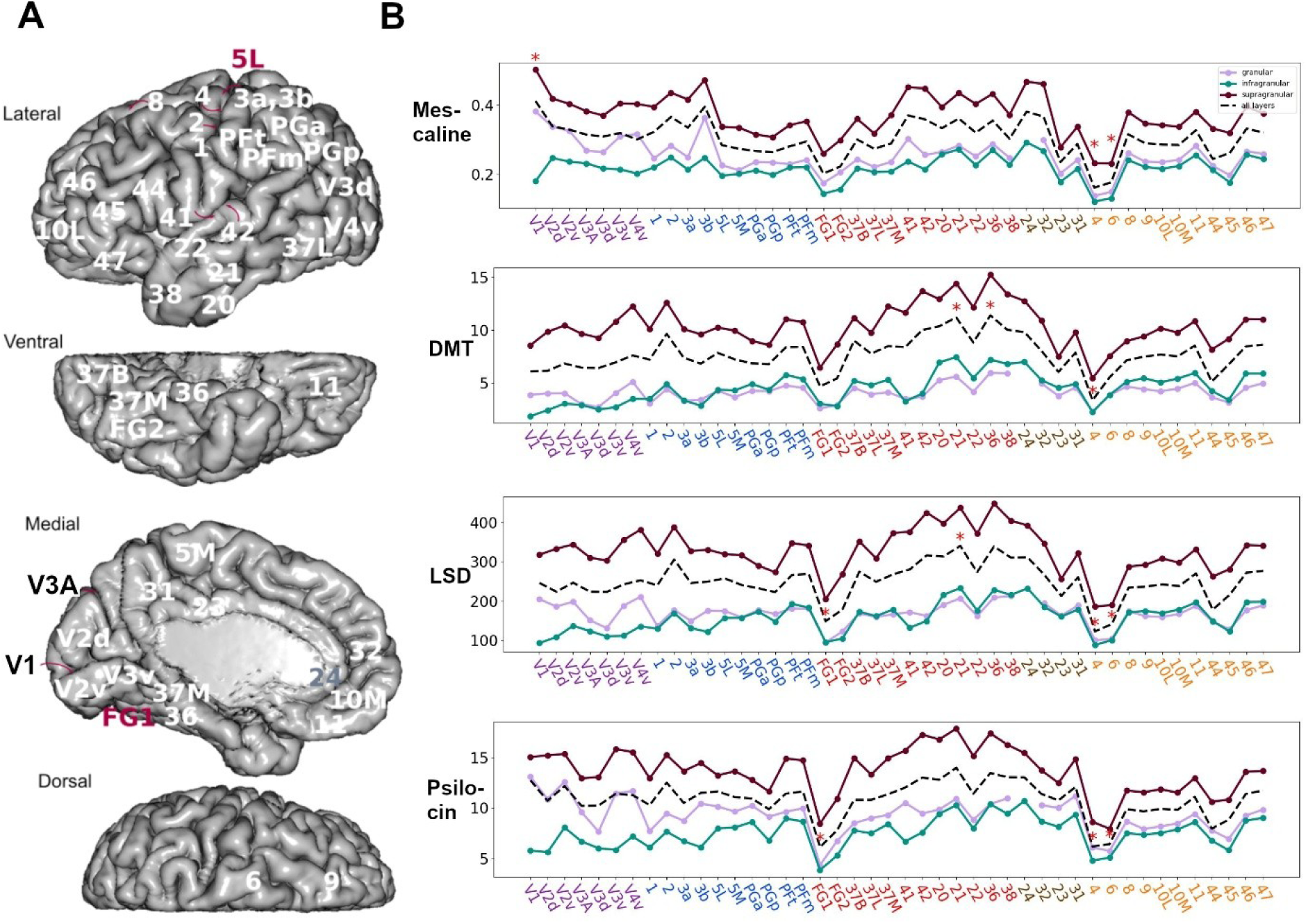
Autoradiography-derived drug distribution profiles of classic hallucinogens. **A)** Display of the analysed cortical areas on the brain’s surface, adapted from [22] which is licensed under CC-BY 4.0. **B)** Layer-resoluted distribution profiles of classic hallucinogens across cortical areas for Mescaline, LSD, DMT and Psilocine. The coloured lines indicate the action strengths within cortical layers as per the legend, the dashed line indicates action strength across the whole cortical surface per area. The colour-coding of x-axis labels indicates occipital (purple), parietal (blue), temporal (red), cingular (brown) or frontal (gold) location. Red asterisks indicate outliers as determined by z-scores > 1.96

### Subcortical drug distribution profiles

To expand the perspective on rapid antidepressant action beyond regional and laminar cortical characteristics, we next turned to subcortical structures. Using PET-derived neurotransmitter receptor density maps, we calculated bilateral drug distribution profiles of classic hallucinogens in the accumbens nucleus, caudate nucleus, putamen, pallidal globe, amygdala, hippocampus and thalamus (**Figure 3A, B**). Akin to the cortex, classic hallucinogens were heterogeneously distributed throughout subcortical structures, but largely interhemispherically symmetric. Generally weak values were found in caudate nucleus, pallidal globe and thalamus, and generally stronger values in amygdala and hippocampus in all substances.

**Figure 3.**
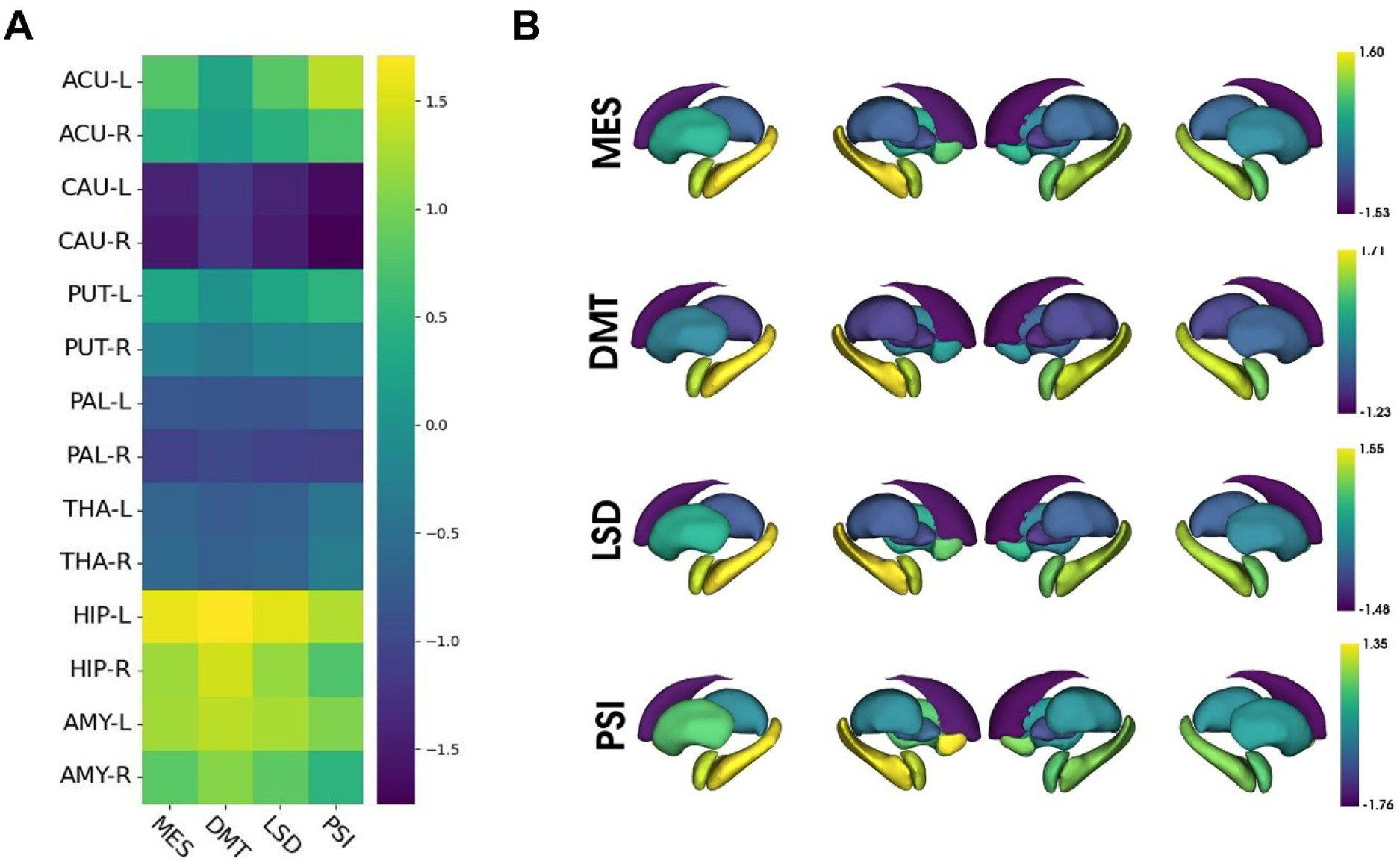
PET-derived subcortical classic hallucinogen drug distribution profiles. **A)** Drug distribution profiles in subcortical structures. **B)** Subcortical projections of drug action values of Mescaline (MES), LSD, DMT and Psilocin (PSI).

### Emphasis on Ketamine

Finally, we investigated the drug distribution profiles of the RAAD Ketamine, with a specific emphasis on exploring the influence that putative 5-HT2a and D2 affinities have on the drug’s distribution profile. To this end, we generated two distinct drug distribution profiles as in the previous analyses: In one set of drug distribution profiles, only the NMDA affinity was considered, while in the other one, the binding to high-affinity subtypes of 5-HT2a and D2 receptors as per [16] were additionally incorporated. The PET-derived NMDA-only cortical drug distribution profile was more homogeneous in comparison to the one additionally considering high-affinity subtypes (**Fig. 4A, C**). In the neuroanatomical decoding, trends towards patterns apparent in drug distribution profiles of classic hallucinogens are apparent in NMDA-only drug distribution profiles, such as moderately higher values in the limbic network, paralimbic and dysgranular cortices, which get more pronounced when considering also high-affinity subtype affinities (**Fig. 4C**). The autoradiography-derived drug distribution profiles show only minimal deviations from one another. Drug action values are consistently highest in the supragranular layer across areas. Across areas, visual cortices show high relative action values, while low relative values are found in isocortical prefrontal and premotor cortices (**Fig. 4B**). In subcortical areas, Ketamine drug distribution profiles show high values in putamen and low values in hippocampus and amygdala, a trend that is more pronounced when considering high-affinity subtype affinities (**Fig. 4D**)

**Figure 4.**
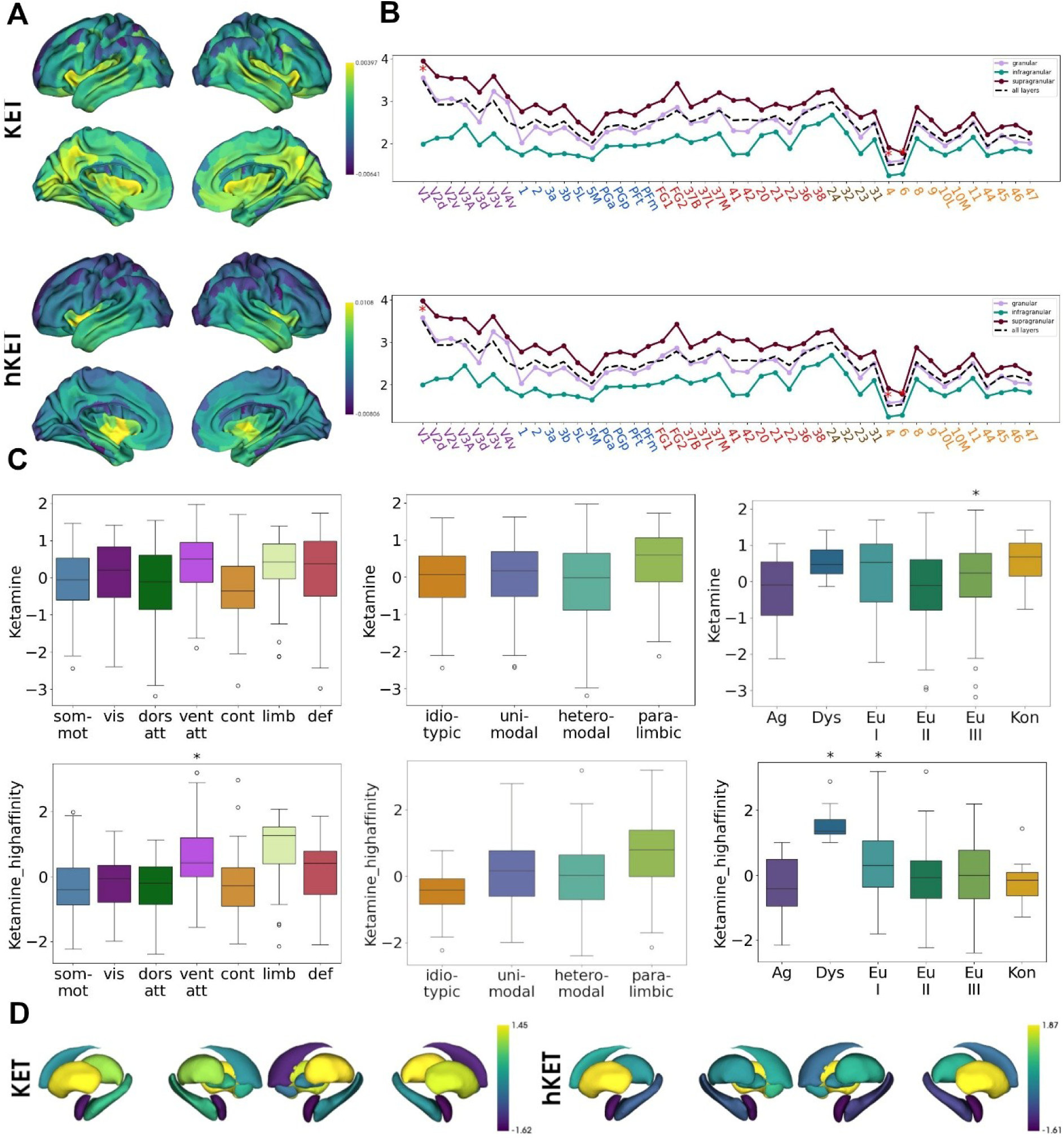
Influence of 5-HT2a and D2 high affinity subtypes on Ketamine drug distribution profiles. **A)** Cortical projection of PET-derived Ketamine drug distribution profiles, without (KET) (top) and with (hKET) (bottom) consideration of high affinity subtype affinities. **B)** Ketamine drug action strengths derived from autoradiography data across cortical areas, without (top) and with (bottom) consideration of high affinity subtype affinities. **C)** Neuroanatomical decoding of PET-derived Ketamine drug distribution profiles analogous to Figure 1B, without (top) and with (bottom) consideration of high affinity subtype affinities. **D)** Subcortical projection of PET-derived Ketamine drug distribution profiles, without (left) and with (right) consideration of high affinity subtype affinities.

### Stability analyses

To exclude parcellation effects, we repeated our cortical PET-based analyses using different parcellation granularities, across which they remain stable (**Supplementary Figures 1-5**).

## Discussion

In this work, we dissect the group of rapid-acting antidepressants, of which we survey classic hallucinogens and Ketamine, from a functional anatomical perspective. Through leveraging pharmacodynamic affinity profiles and the human brain’s chemoarchitecture, we calculate drug distribution profiles in cortical and subcortical areas that give a comparative measure indicating how strong a drug would act in a region of interest. PET-based drug distribution profiles revealed that classic hallucinogens act strongly in association cortices, especially in the temporal lobe and the insula, and generally show their strongest drug action values in dysgranular cortices of limbic functionality. In the subcortex, they show high values in the amygdala, hippocampus and nucleus accumbens. Similar trends in drug distribution patterns are apparent when purely considering the NMDA affinity of Ketamine, which become more pronounced when incorporating putative binding to high-affinity subtypes of the 5-HT2a and D2 receptors. Finally, autoradiography-based drug action profiles indicate strong action of classic hallucinogens in the multimodal Brodmann areas 21 and 36 in the temporal lobe, and within the supragranular cortical layer across the whole cortex, while Ketamine generally shows a stronger trend towards visual areas.

First, we investigate RAAD drug distribution profiles across the whole brain. Our results show that RAAD drug action strengths are heterogeneously distributed across the human cerebral cortex. Classic hallucinogens show low drug action values in the paracentral lobule and at the occipital pole, and high drug action values in the temporal lobe and the insular region. The neuroanatomical contextualisation analyses allow us to derive more detailed implications from the distribution patterns. Our results suggest that classic hallucinogens’ drug action strengths vary as a function of cognitive processing complexity. This is apparent in the clear alignments of the lower drug action values with the somato-motor network, and of the higher drug action values with the limbic and default-mode networks in the functional network decoding analysis. Limbic network and DMN are maximally different from the somato-motor network with regards to their functional connectivity characteristics, as shown by their opposite locations on a principal gradient of functional connectivity [28]. A potential dependence of the heterogeneous distribution of relative action strengths on functional aspects is similarly reflected in the analysis of cytoarchitectural classes, where it increases along a gradient of cognitive processing complexity [25]. A more detailed view on the relationship to histological characteristics is provided through the decoding analysis that employs laminar architecture-derived cortical types, where classic hallucinogens show their strongest relative activity values in dysgranular cortices, which act as a site of functional and structural mediation between evolutionarily older agranular and more recent eulaminate areas [29].

Extending our models towards a layer-specific activity model, we find that, across all cortical areas investigated, RAAD show their strongest drug action values in the supragranular layer. Cortical layering is an important determinant of cytoarchitectural composition and constitutes the foundation for neuroanatomical cartography studies that run orthogonal to the well-known field of studies that parcellates the cortex horizontally upon inter-areal boundary characteristics [30]. Cortical layers also differ with regards to their connectivity profiles and supposed functionality. Generally, deep layers receive efferent fibers from cortical regions with a lower myelination profile and send afferent fibers to cortices with a higher myelination profile and thalamic nuclei. Analogously, superficial layers receive efferent fibers from cortical regions with a lower myelination profile, and from hippocampus and amygdala, and send afferent fibers to cortices with higher myelination profiles [31]. Regarding their functionality, the superficial layers are hypothesised to perform the bulk of cortical signal integration and computation, combining the inputs from previous cortical areas, while deep layers relay the outputs of these computations to subsequent cortical areas or thalamic nuclei [32]. Our analysis showed no clear relationship between the complexity of functionality realised in cortical areas and the between-layer distribution of RAAD action strengths. From the within-layer functional perspective, however, our results suggest that RAAD primarily influence signal integration and computation that takes place in the supragranular layer, which is consistent with findings that supragranular-located 5-HT2a-expressing GABAergic interneurons show increased transcription rates after 5-HT2a agonism in a rodent model [33,34]. The high supragranular drug action values furthermore support theories that designate deep-layer pyramidal cells as important targets for RAAD [13,15], as their 5-HT2a-carrying apical dendrites are located in supragranular layers.

Moreover, our autoradiography-derived drug distribution profiles establish further connections of pharmacological action to functional anatomy. First, it is important to highlight a general correspondence between the autoradiography-derived and the PET-derived drug distribution profiles, as both locate high drug action values within the temporal lobe, and here, especially within multimodal cortices that are generally considered (para)limbic structures, such as the perirhinal cortex (Area 36) and the temporal pole (Area 38). A strong example of classic hallucinogens’ high affinity to limbic structures can be seen in direct comparison of cytoarchitectural areas in the cingulate gyrus. Here, the periarchicortical area 24, which is thought the be primarily involved in emotional processing, shows markedly higher drug action values than the isocortical area 32, which is of more general higher-order cognitive processing functionality [35]. The functional relevance of areas with high drug action values for emotional processing is also apparent in the aforementioned temporal limbic areas [36,37]. That the tie to emotional processing also extends beyond limbic structures is apparent when comparing the lower-valued fusiform gyrus (FG) with the higher-valued superior temporal gyrus (Area 22). While both areas are involved in high-level processing of visual stimuli of faces, FG is more relevant for the recognition of static features of the face, while the superior temporal gyrus contributes more to recognising dynamic and changing features, such as emotionally-laden facial expressions [38]. A tie to alterations in affective stimulus processing was similarly proposed for the multimodal temporal area 21, a further site of strong drug action values, given its involvement in depression-associated circuit alterations [39].

Our analysis of subcortical structures can put the relationship between classic hallucinogens’ drug action strengths and emotional processing into a wider context. Here, we see that hippocampus and amygdala, as important limbic structures, showed the highest drug action values. Tracer-based connectivity studies show that amygdala receives input from higher-order sensory and limbic association cortices, especially areas 24 and 38, at the same sites from which it projects to prefrontal cortices [36]. Furthermore, amygdala projects to the perirhinal cortex, area 36, which in turn projects towards the hippocampus via entorhinal relays [40]. Importantly, all listed feed-forward connections are mirrored by feed-back projections, creating a network between structures with high drug action strengths of classic hallucinogens that is heavily implied in emotion processing and formation of emotional memories [41].

We can use a synopsis across modalities to discuss our results within the wider context of hypothesised modes of action of rapid-acting antidepressants. Our results imply that rapid-acting antidepressant’s main sites of action are (para)limbic cortical and subcortical structures. In this regard, our findings differ at first glance from the anatomical and functional claims of the REBUS model, which asserts that classic hallucinogens lessen the influence of highest-level beliefs through direct action in the Default Mode Network. There is, however, strong evidence for relevant alterations of limbic and DMN interplay through psychedelic drugs that might help reconcile this difference. Functional imaging studies show discordant BOLD signal changes in limbic versus other higher-order cortices after the administration of psilocybin [42], consistent with reduced functional coupling between hippocampus and parahippocampal cortices and cortical nodes of the DMN under psilocybin [43]. Reduced functional interplay between DMN and limbic structures could be specifically associated with the psychedelic effect of ego-dissolution [44]. Our results therefore add to the body of evidence that suggest alterations of functional connectivity patterns in DMN, which can indeed be observed after application of classic hallucinogens [45], are a secondary effect that is mediated through drug action in (para)limbic structures, in which RAAD might produce an increased repertoire of connectivity motifs as a primary effect [43]. Summarised, our study highlights the role of regions that are functionally involved in emotion processing, and act as intermediaries between prefrontal associative cortices and higher-order sensory cortices, in rapid antidepressant action.

In a final analysis, we investigated the influence that affinities of Ketamine to high-affinity subtypes of 5-HT2a and D2 receptors have on drug distribution profiles. While Ketamine is a clinically efficacious rapid antidepressant drug, it is primarily viewed as an NMDA antagonist, challenging serotonergic conceptions of rapid-antidepressant action [13] and spawning its own theories [46]. Our results show that incorporating 5-HT2a and D2 affinities yields more heterogeneous cortical PET-derived drug distribution profiles which share hallmark features of classic hallucinogens, such as the significant alignment to dysgranular cortices and a clearer differentiation along the sensory-fugal gradient. In the subcortical drug action profiles, Ketamine shows high values in putamen and nucleus accumbens, rather than in hippocampus and amygdala, where classic hallucinogens show high values. Regarding the autoradiography-derived relative drug action profiles, however, Ketamine shows more dissimilarities to classic hallucinogens. While they share that the supragranular layer consistently shows the highest values, the trend towards high values in limbic structures is not as pronounced, with high values rather being found in early visual cortices. Importantly, incorporating high-affinity subtype affinities for 5-HT2a and D2 only resulted in minuscule alterations in the drug’s relative action profile. There are two possible explanations for this difference in drug alteration profiles using either PET-derived or autoradiography-derived neurotransmitter receptor maps: First, as no measurements of D2 distributions was present in the autoradiography dataset, this affinity could not be considered in generating the layer-resoluted drug action profiles. Second, relative drug action profiles from autoradiography were calculated from absolute receptor densities (in fmol/mg protein). One constraint of PET, however, is that it yields not absolute, but relative protein densities [47], which might lead to over-estimations of contributions of low-density receptors, and under-estimation of contributions of high-density receptors specifically in PET-derived relative drug action profiles. This fact is of lesser importance in classic hallucinogens, which are primarily 5-HT1a and 5-HT2a-affine, as these receptors show a comparable absolute range of densities [23,48], but skews in cortical PET-derived relative drug action profiles for high-affinity Ketamine towards 5-HT2a and D2 distributions might be more pronounced, since there is evidence for NMDA absolute densities being relevantly higher in the human cerebral cortex [23,49]. Regarding the relevance of Ketamine’s action at high-affinity subtypes of 5-HT2a and D2 receptors, our results therefore paint an inconsistent picture, and, at best, offer weak support for the importance of direct serotonergic signaling. Yet it is unclear whether high-affinity subtypes are evenly distributed, which our method assumes. Further insights into this question could be gained through studies that co-administer 5-HT2a antagonists and Ketamine, as has been done to test the receptor’s relevance for the effects of classic hallucinogens [50].

In sum, our work highlights the role of brain areas that are functionally involved in the processing and regulation of emotions in rapid antidepressant action, adding important nuances to previously proposed mechanisms and suggesting alternative, but coherent, interpretations of functional imaging findings [13,43]. Our insights support novel efforts that focus on treatment targets outside of the frontal lobe, such as the insula [51], and underscore the importance of the brain’s chemoarchitectural organisation in understanding intricate structure-function relationships [18,19].

## Materials & Methods

### Affinity profile generation

To generate the molecular affinity profiles, previously published inhibition constants were collected from multiple sources. Ki values for classic hallucinogens were drawn from [21]. For Ketamine, Ki values for high-affinity state 5-HT2a and D2 receptors were drawn from [16]. Ki values were converted to nanomolar. Ki values were capped at 100000, which was also used to represent missing data.

### PET data preparation

To assess neurotransmitter receptor densities in the brain as a prerequisite for generating cerebral antidepressant binding profiles, we leveraged a collection of open-access volumetric PET maps from healthy participants [19]. For cortical receptor densities, we performed PET data preparation similar to a previous description [18] - briefly, the PET density maps were parcellated based on the Schaefer parcellation [52]. Following original author recommendations, we extracted data from surface-projected receptor density maps for 5-HT1a and 5-HT2a [48]. Granularities from 100 to 400 parcels were used to replicate the results in order to control for parcellation-based effects. Subcortical receptor densities were extracted bilaterally for accumbens and caudate nuclei, pallidum, pallidal globe, thalamus, amygdala and hippocampus as averages per structure. The resulting parcel-wise intensity values were z-score normalized per tracer. Both processing steps were performed separately for the cortical and subcortical compartments. Density maps of the following receptors as quantified through PET scans were used: 5-HT1a [48,53], 5-HT2a [48], NMDA [54], D1 [55] and D2 [56].

### Autoradiography data preparation

The autoradiography data were collected as described in [23], for which we refer for a detailed description. In summary, receptor densities were quantified by in vitro receptor autoradiographical analysis, using an incubation protocol of post-mortem brain slices with tritium-labeled ligands, subsequent exposure to tritium-sensitive films and densitometric analysis. Four hemispheres stemming from three brains of donors with no record of neurological and psychiatric diseases were used. The specific anonymized autoradiography receptor density data was published accompanying [22]. In this study, we used the autoradiographic density profiles of the following receptors: 5-HT1a, 5-HT2, alpha1, alpha2, D1, NMDA.

### Drug distribution profile generation

To calculate the anatomical drug distribution profiles, the following receptors were used to calculate the cortical and subcortical antidepressant binding profiles. For each substance, a unit-less value a per region was calculated as

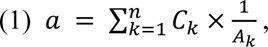

with C and A being one-dimensional arrays representing the same receptor density (*C*) and affinity (*A*) at location *k*, and *n* representing the total number of receptor densities as quantified by PET or autoradiography measurement, respectively. For each region of interest, the value *a* represent the expected strength of drug action based on that region’s chemoarchitecture and the affinity profile of the drug in question.

### Functional network and cytoarchitectural classes decoding

To contextualize PET-derived cortical RAAD drug distribution profiles, we leveraged a resting-state fMRI-based partition of the cortex into seven functional networks [26], a theoretical framework of cytoarchitectural classes that describe a sensory-fugal gradient of cognitive processing complexity [25], and laminar architecture-derived cytoarchitectural classes [24,31,57].

### Null models and statistical significance testing

To assess the significance of alignment of cortical drug distribution profiles with neuroanatomical measures, we applied spin permutation [58] to generate randomly permuted brain maps by random-angle spherical rotation of surface-projected data points, which preserves spatial autocorrelation, using the brainspace python package [59] (https://github.com/MICA-MNI/BrainSpace). Vertex values that were rotated into the medial wall, and values from the medial wall that were rotated to the cortical surface, were discarded [60]. In each approach, we generated 1000 permuted brain maps. Alignment was determined as significant when true mean values within the respective regions were either below the 5th or above the 95th percentile of the distribution of randomly permuted values.

## Ethics declaration

This study re-analysed previously published open-access data. No subject-level data was used. It was approved by the ethics committee of the medical faculty, University of Tübingen, Germany (Project Number 492/2024BO2).

## Conflict of interest statement

The authors declare no conflict of interest.

## Acknowledgements

B.H. acknowledges funding from the Max Planck School of Cognition’s Clinician Scientist programme.

T.K. acknowledges funding from the Research Council of Norway (#323961) and the European Research Council (ERC CoG, #101086793, HealthyMom). S.L.V. acknowledges funding from the Max Planck Society (Lise Meitner excellence programme), the Jacobs Foundation, the Hector Fellow Academy (Research Career Development Award) and the Helmholtz International BigBrain Analytics and Learning Laboratory (HIBALL).

## Code and data availability

PET-derived neurotransmitter receptor maps can be accessed through https://netneurolab.github.io/neuromaps/. The specific anonymized autoradiography receptor density data can be accessed at https://github.com/AlGoulas/receptor_principles/. The code and other resources to reproduce the main figure results is available at https://github.com/bhaenisch/nchem_raad/.

## Supplementary figures

**Supplementary Figure 1.**
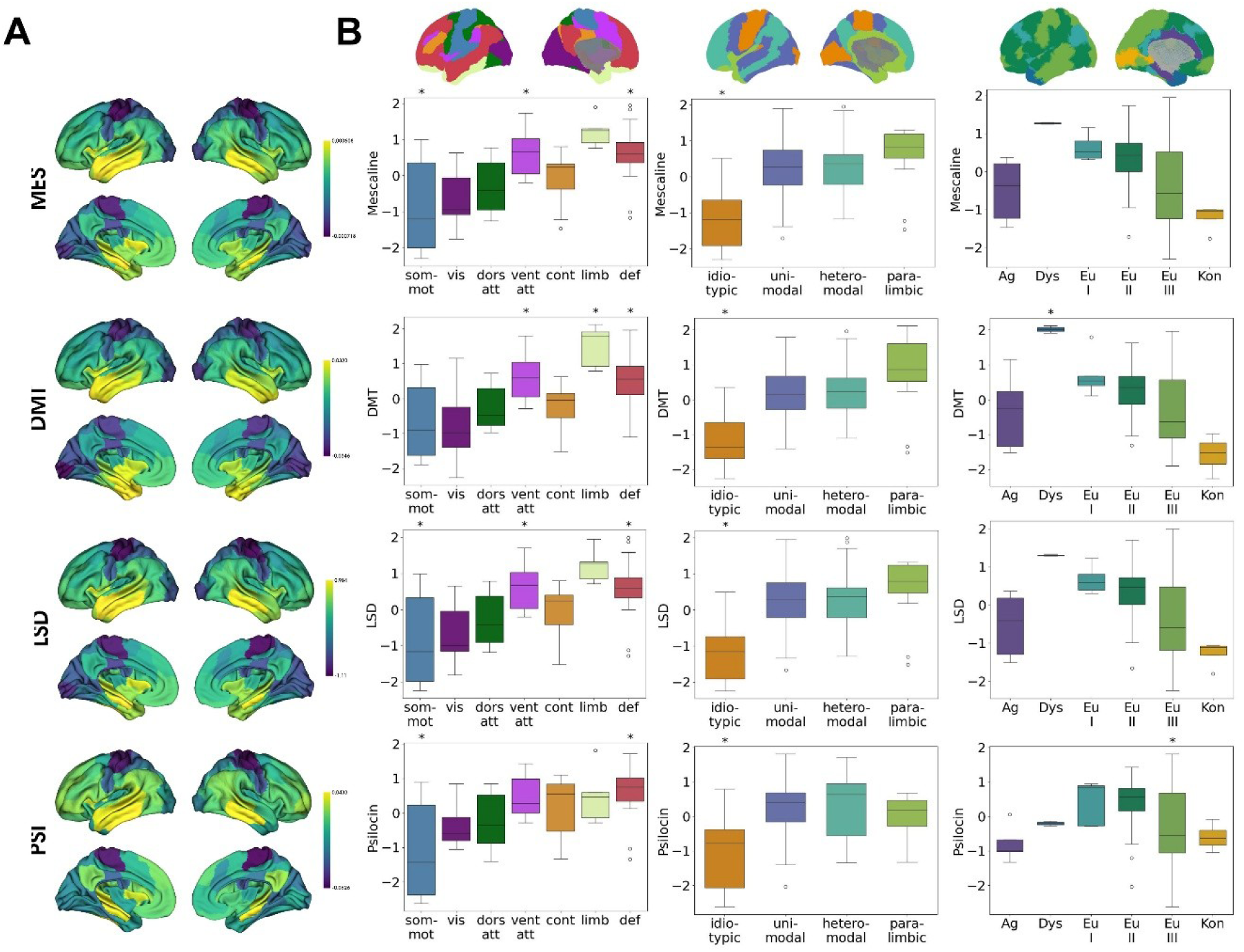
Replication of Figure 1 using a granularity of 100 Schaefer parcels.

**Supplementary Figure 2.**
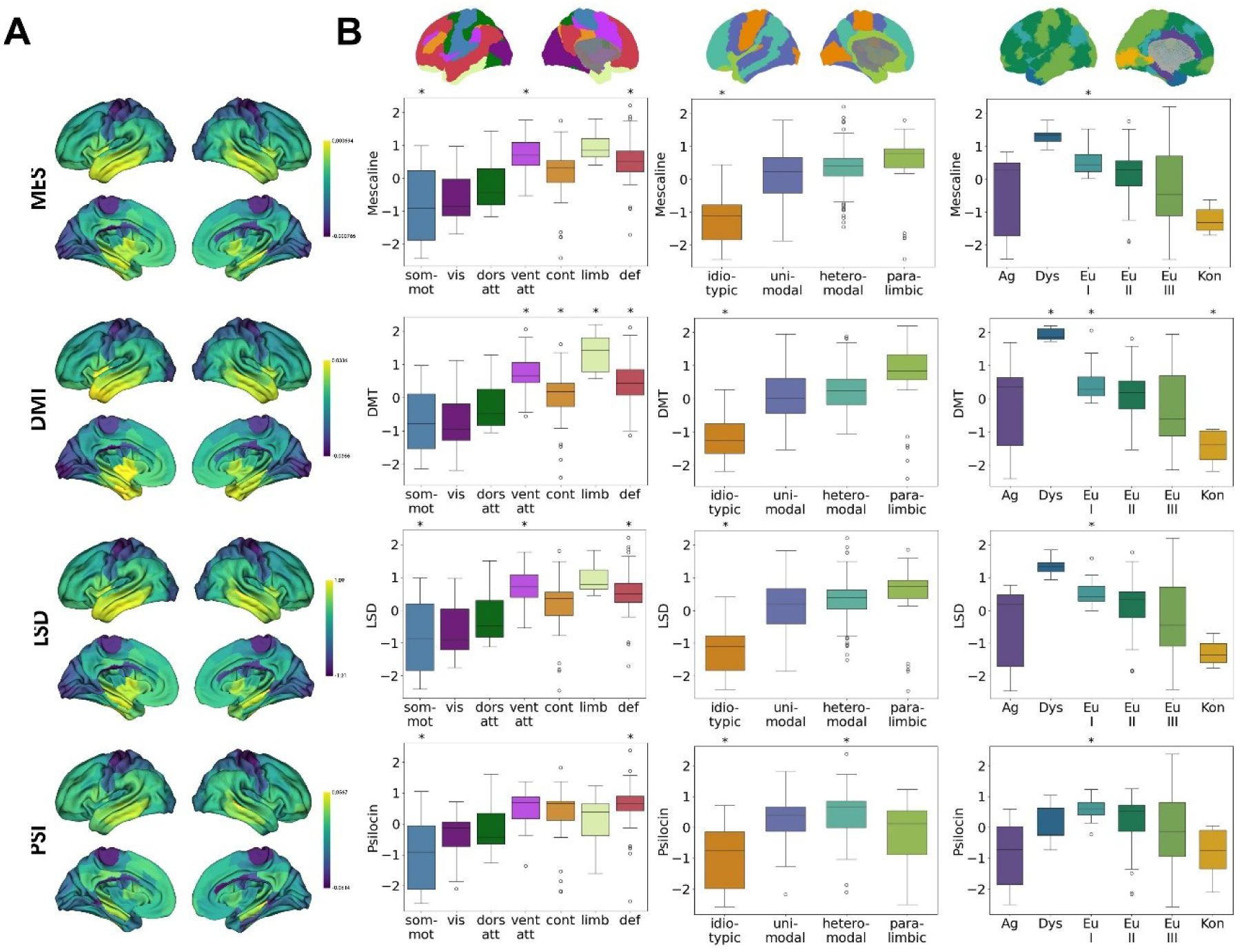
Replication of Figure 1 using a granularity of 200 Schaefer parcels.

**Supplementary Figure 3.**
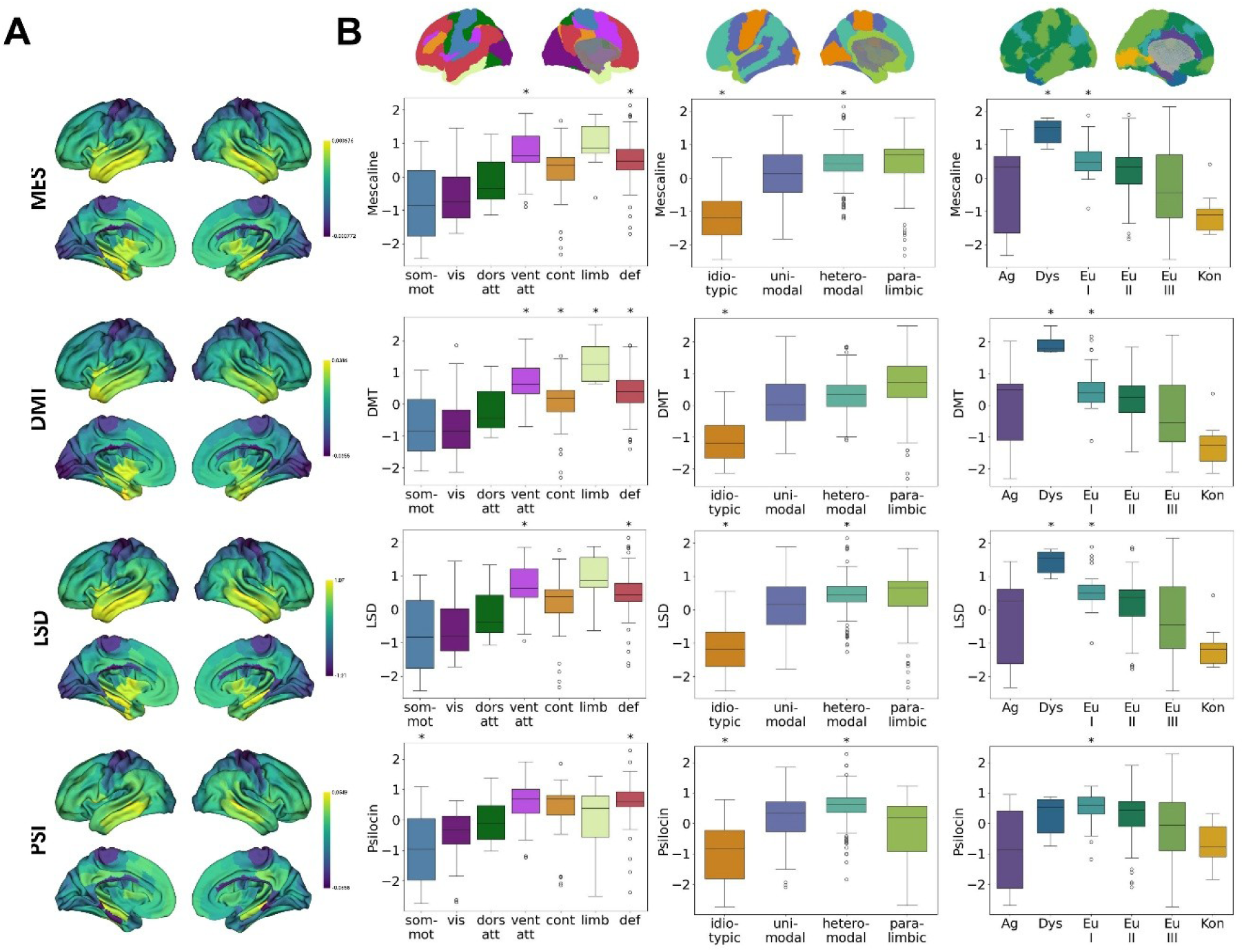
Replication of Figure 1 using a granularity of 300 Schaefer parcels.

**Supplementary Figure 4.**
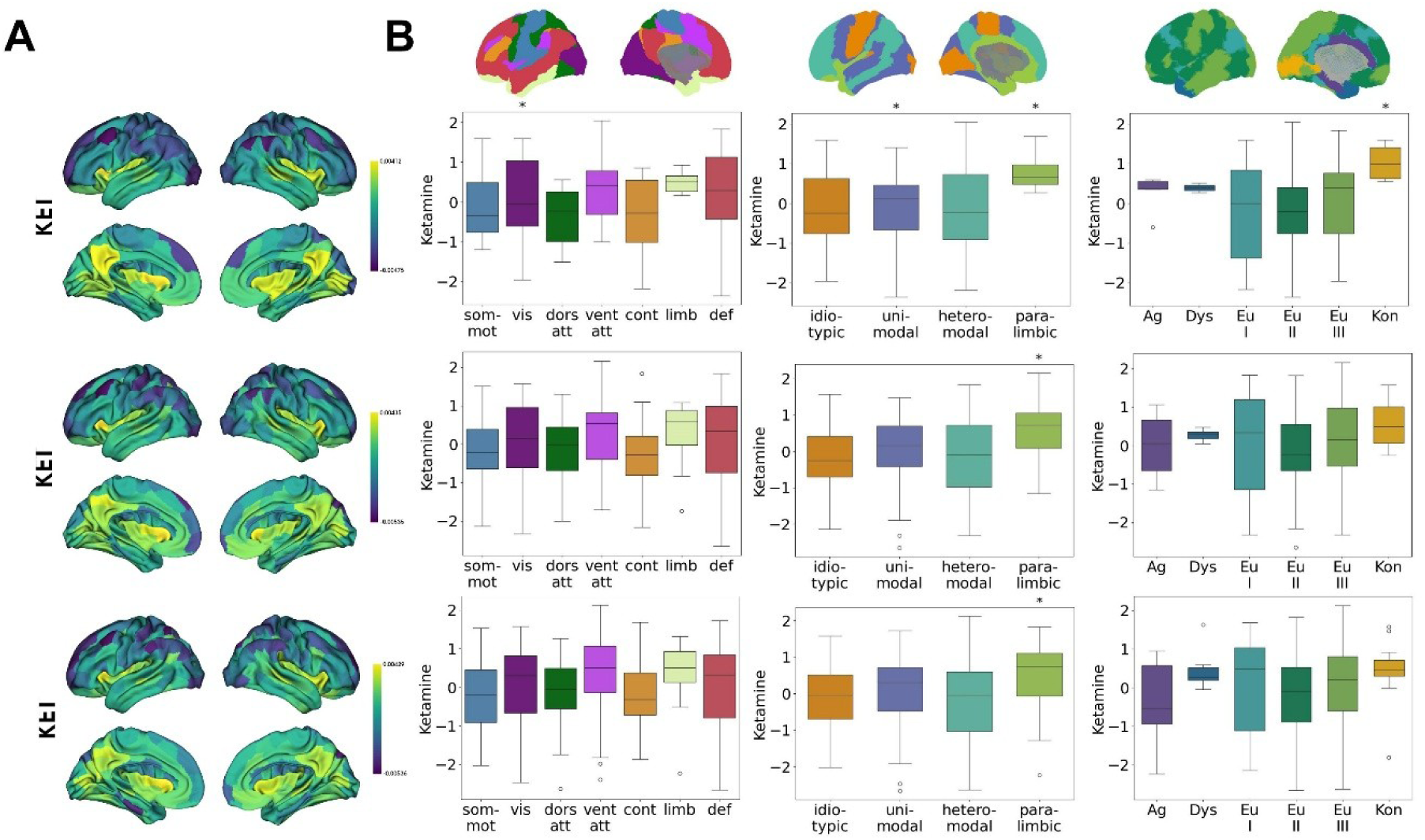
Stability analysis of cortical PET-based Ketamine drug distribution profiles (NMDA-affinities only) in granularities of 100 (top), 200 (middle) and 300 (bottom) Schaefer parcels.

**Supplementary Figure 5.**
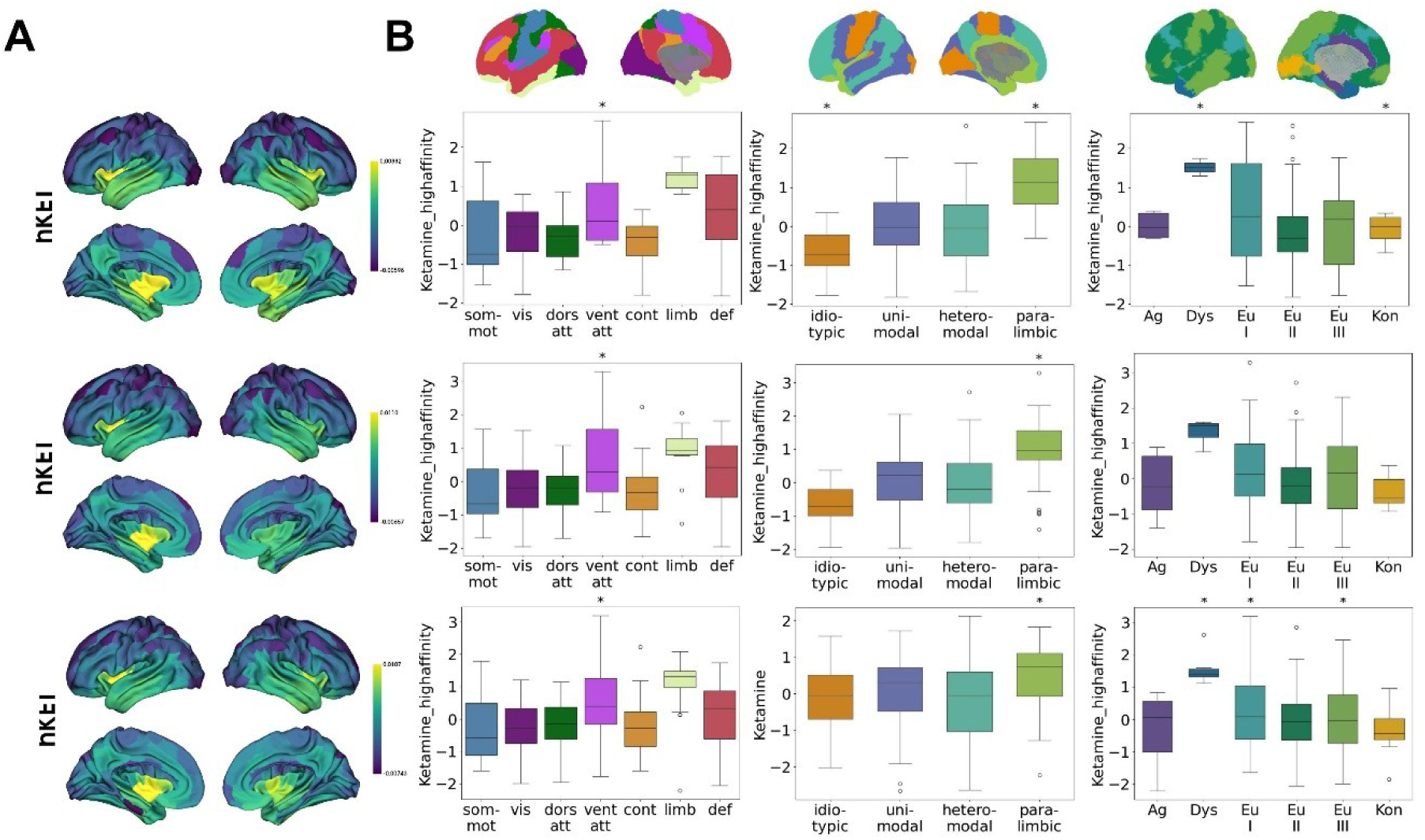
Stability analysis of cortical PET-based Ketamine drug distribution profiles (NMDA, 5-HT2a and D2 affinities) in granularities of 100 (top), 200 (middle) and 300 (bottom) Schaefer parcels.

## Notes

### Competing Interest Statement

The authors have declared no competing interest.

